# MaxCAT-Net: Hybrid transformer-based skin lesion classification and lightweight segmentation with calibration and cross-dataset validation

**DOI:** 10.64898/2026.11.01.763389

**Authors:** Sadek Al Prince, Israt Fatema Shorna

## Abstract

Class imbalance, poor calibration of confidence and poor generalisation under different imaging conditions are restrictions of automated analysis of dermoscopic skin lesions. We propose a hybrid classifier MaxCAT-Net (MaxViT + CBAM + Attention Transformer), which consists of a pre-trained MaxViT-Tiny backbone and the combination of CBAM and two-layer lightweight Transformer encoder and evaluate four encoder-decoder networks for lesion segmentation as a separate task on ISIC 2018 Task 1. All the images of exact/near duplicate were removed and the remaining 10,012 images were distributed lesion-wise (72/14/14) without two or more images having the same type of lesion in HAM10000. On the test set (*n* = 1,405), MaxCAT-Net outperformed the baselines of ResNet152-CBAM, MobileNetV3-CBAM, GhostNet-hybrid and single MaxViT-Tiny-CBAM with an accuracy of 0.869 (95% CI 0.847–0.890), balanced accuracy of 0.742, macro-F1 of 0.751 and macro-AUC of 0.944, although ablation gains in macro-F1 (0.017–0.022) were not statistically conclusive. By scaling the temperature, the ECE was decreased from 0.099 to 0.038. In the case of the PAD-UFES-20 dataset, however, the accuracy fell to 0.341 and the melanoma recall rate to 0.096 while for the HAM10000 dataset, the melanoma recall rate was only 0.587. A Swin-T-encoder U-Net was able to achieve a Dice score of 0.903 in segmentation (520 test images), whereas a FasterNet-T0 U-Net (1/10th of the number of parameters) had a Dice score of 0.901 (Holm adjusted *p* < 0.002), and a standard U-Net had a significantly lower Dice score of 0.888. All results are based on one seed and show trends—not definite improvement.

## 1 Introduction

Eight out of ten of the most common cancer in the world is skin cancer and the most common form of skin cancer that causes skin cancer deaths is melanoma. In 2025 it is estimated that 104,960 new cases of invasive melanoma will be diagnosed in the U.S. and 8,430 people will die of the disease.[1] The prognosis is strongly correlated with the stage of diagnosis, early and correct diagnosis are crucial. Until now, diagnosis has been based on subjective assessment and dermoscopy that in daily practice depends on the experts’ experience and skill. There is considerable observation to observation variation and dermatologists are not widely available.

Thus deep learning is emerging as an important technique in the domain of Computer Aided Diagnosis (CAD) in dermatology. In a study conducted by Esteva et al., they showed a CNN training with 129,450 clinical images, can classify the skin lesions at a level comparable to board certified dermatologists.[2] Similarly, Haenssle et al., collected the results of 58 dermatologists in recognising dermoscopic melanoma.[3] Due to the release of large public databases like HAM10000 (10,015 dermoscopic images for 7 classes)[4] or ISIC challenge datasets[5, 6] and BCN20000[7] progress in the field has been improved.

Here are the three task-formulations which are predominant in this literature. Classification involves giving a lesion image a diagnostic classification. The detection takes place in a single shot and the lesions are localised using bounding boxes and labelled. Segmentation (cutting the lesion off from other skin). The tasks are supportive of each other. Clinically meaningful descriptors (such as asymmetry, border irregularity, diameter) can be obtained by proper definition of boundaries and restricting the model to only the inner part of the lesion can improve model robustness and interpretability of classification.[8] Even segmentation is difficult because of the presence of hair, air bubble, ruler marks, low contrast with the skin, fuzzy boundary and varying acquisition conditions.[8]

Current trend in architecture has been changed from classic U-Net[9] and its variants such as UNet++,[10] Attention U-Net[11] and nnU-Net[12] to Transformer based or hybrid CNN-Transformer such as TransUNet[13] and Swin-Unet[14] based on ViT[15] and Swin Transformer[16] respectively. Moreover, efforts have been made to adapt the model for use with resource-constrained devices like UNeXt[17] and EGE-UNet[18] and the recent real-time detection paradigm (YOLOv10[19]) has been adapted for skin cancer detection on dermoscopic images.[20]

But there is still room for improvement: the generalization of skin tone to imaging device, the variability of performance between skin tone[21, 22] and the lack of transparency and trustworthy prediction are still the limiting factors. In this paper, we propose to use the MaxViT-Tiny backbone[23] and Convolutional Block Attention Module (CBAM)[24] and lightweight Transformer encoder (MaxCAT-Net: MaxViT + CBAM + Attention Transformer) to address these issues and perform seven-class dermoscopic skin-lesion classification under stringent leakage-controlled and lesion-wise protocol. Besides, there are four encoder-decoder networks that assess as a stand-alone task for lesion segmentation on ISIC 2018 Task 1.[5] The contributions are mainly:

- All the MaxCAT-Net (a hybrid MaxViT-Tiny classifier + CBAM + mini-Transformer) and four-variant ablation (backbone only, +CBAM, +Transformer, +both) are trained with the same split and hyperparameters and their random seed.
- Then, a leakage-controlled evaluation pipeline (duplicate removal, lesion-wise stratified splitting, perceptual-hash audit of test-to-train near-duplicates, and lesion-cluster bootstrap confidence intervals) is presented, and compared to ResNet152-CBAM, MobileNetV3-CBAM, a GhostNet hybrid, and a single MaxViT-Tiny-CBAM baseline on the same split.
- Calibration (Temperature Scaling) analysis, Selective Prediction and Cost of Computation (ONNX conversion and INT8 quantisation).
- Comparing the four networks (standard U-Net, Attention U-Net, U-Net with FasterNet-T0[25] and U-Net with Swin-T) in terms of accuracy and computation cost related to ISIC 2018 Task 1.
- Cross-dataset evaluation (on PAD-UFES-20[26]) and skin-tone related subgroup analysis and qualitative explainability analysis (Grad-CAM++ and Occlusion Maps).

## 2 Literature Review

### 2.1 Skin Lesion Classification

Initial CAD systems have relied on the handcrafted colour and texture features, and classical classifiers. They were subsequently replaced by deep CNNs by Esteva et al.[2] and subsequent studies showed results that were either as good as or better than dermatologists in controlled reader studies.[3] The usage of transfer learning using the backbones of EfficientNet[27] and ConvNeXt[28] and Vision/Swin Transformers[15, 16] have been used later. Ensembles were introduced and both class-imbalance handling and augmentation were done on HAM10000[4] and ISIC challenges[5] frequently. Although human-computer collaboration research showed that AI assistance is beneficial to clinicians, this is dependent on readers’ experience and AI reliability.[29]

### 2.2 Detection-Based Approaches

Real-time object detector can be used as the screening workflow as they can localise and classify in one forward pass. YOLOv10 introduced an end-to-end design which took away the non-maximum suppression at the time of inference which increased the speed-accuracy trade-off.[19] The next one was developed by Prince et al.[20] with computer vision system for skin cancer detection from dermatoscopic image using YOLOv10 in ICECTE 2026 conference, which is classified into multi-class. This kind of detection based method, apart from the class label, is able to output the location of the lesion but it is in the form of a bounding box and not a pixel level lesion mask.

### 2.3 Skin Lesion Segmentation

A recent survey involving 177 papers discusses the categorization of the input data, model designs (architecture, modules, losses), evaluation and open limitations for analyzing the skin lesion segmentation.[8] Most of the models are encoder-decoder models that are inspired by U-Net.[9] Can be extended to make nested skip connections[10] or attention gates[11] or to a self-configuring pipelines.[12] To achieve this, transformer-based models such as transformer hybrids like TransUNet[13] or Swin-Unet[14] and boundary aware Transformers like FAT-Net[30] can be helpful to have access to long range context and improve the delineation of boundaries. For the purpose of application on mobile or point-of-care, the parameters and complexity of the methods can be reduced, such as the lightweight UNeXt[17] and EGE-UNet.[18]

### 2.4 Joint Segmentation and Classification

Several works are related to both the two tasks, because the location priors provided by the lesion masks during the diagnosis. To simultaneously perform both segmentation and classification, Yang et al.[31] suggested a multi-task network for segmentation and classification. Xie et al. proposed a mutual bootstrapping approach for an iterative improvement of each task in Ref. [32]. These approaches all report that the sharing of representations is beneficial in general to both tasks. This also makes the training much more complicated and usually, test only one data set.

### 2.5 Explainability, Generalisation and Fairness

Fitzpatrick 17k (16,577 clinical images, 114 conditions) was published by Groh et al. and it was shown that the best performance of CNNs is on skin types similar to the training data.[21] Daneshjou et al. (2017)[22] demonstrated that the performance of Dermatology AI is lowered in the case of dark skin tone and rare diseases, using a curated set of diverse clinical images. Rotemberg et al.[33] have shown patient-level contextual information can be useful to identify melanomas. Saliency methods are commonly used for the purpose of interpretability, but not always assessed via lesion masks of clinically relevant features, such as Grad-CAM.[34]

## 3 Methodology

### 3.1 Dataset

#### HAM10000

The experiments are conducted on the HAM10000 dataset[4] available on Kaggle which has 7 diagnostic classes actinic keratoses and intraepithelial carcinoma (akiec), basal cell carcinoma (bcc), benign keratosis-like lesions (bkl), dermatofibroma (df), melanoma (mel), melanocytic nevi (nv) and vascular lesions (vasc). The images were aggregated based on their lesion identifiers (collected from the metadata text file of the HAM10000 dataset) and 7,470 different groups were achieved. The dermoscopic images of two samples of each of seven classes are shown in Fig. 1.

**Figure 1.**
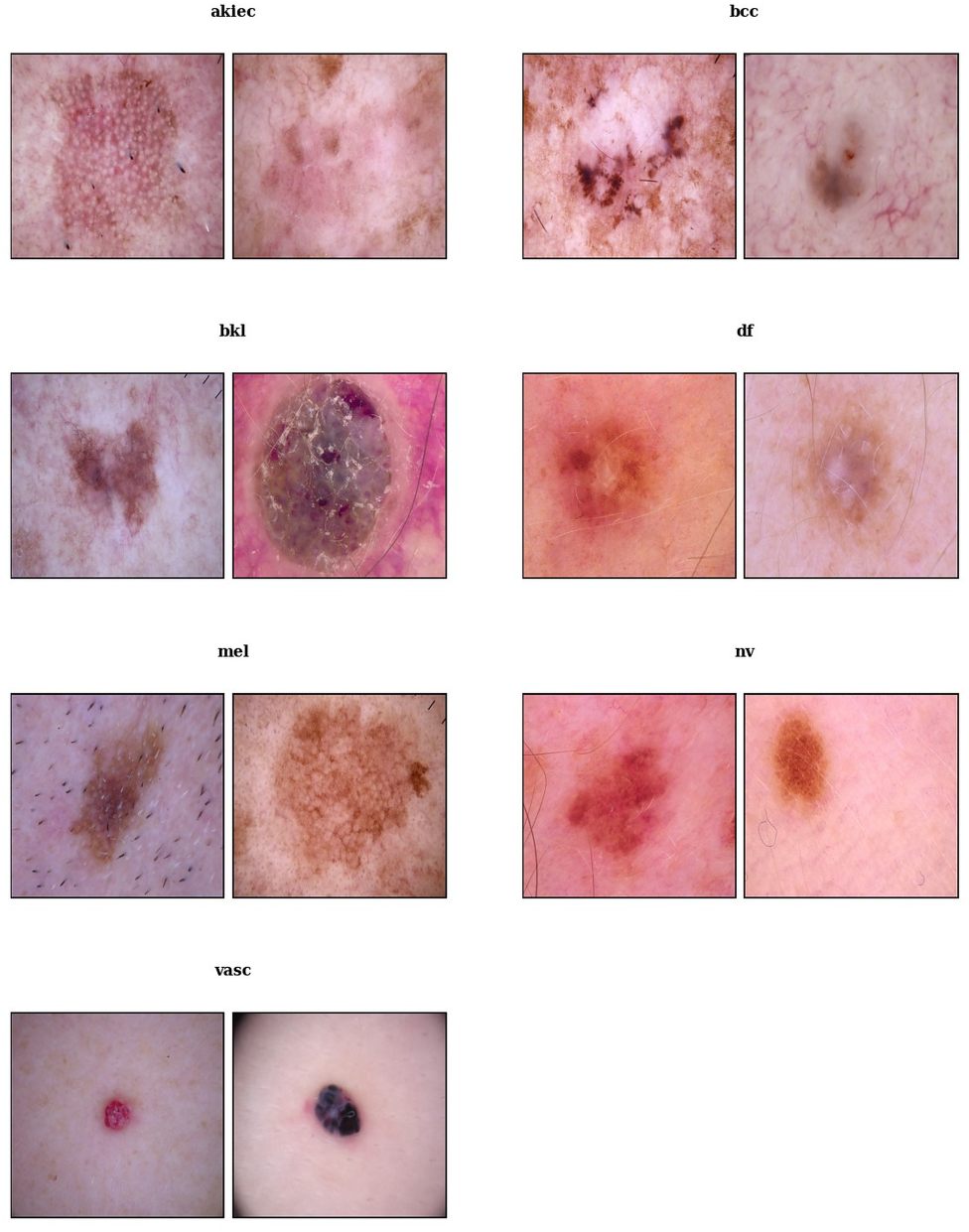
Sample dermoscopic input images from the HAM10000 dataset, showing two representative examples for each of the seven diagnostic classes (akiec, bcc, bkl, df, mel, nv and vasc).

#### Duplicate removal and splitting

All images were verified as not corrupted (no images), of not having the same label but different file name (no images), and of either being exactly the same (2 images removed) or being near-duplicates (1 image removed) using a 16 × 16 difference image hash, resulting in 10,012 images. Next, data was split with a stratified group *k*-fold procedure (*k* = 7, seed = 43), with one fold being used for training, one for validation and one for testing, resulting in an approximately 72/14/14 train/validation/test split. Split is done by lesion identifier so that no lesion is in more than one subset (checked: 0 shared groups). The resulting class distribution is shown in Table 1, and in Fig. 2.

**Table 1:** Class distribution of HAM10000 after de-duplication and lesion-wise splitting.

| **Class** | **Train** | **Validation** | **Test** | **Total** |
|---|---|---|---|---|
| akiec | 238 | 54 | 35 | 327 |
| bcc | 362 | 67 | 85 | 514 |
| bkl | 787 | 165 | 147 | 1,099 |
| df | 85 | 19 | 11 | 115 |
| mel | 782 | 188 | 143 | 1,113 |
| nv | 4,813 | 932 | 957 | 6,702 |
| vasc | 85 | 30 | 27 | 142 |
| Total | 7,152 | 1,455 | 1,405 | 10,012 |

**Figure 2.**
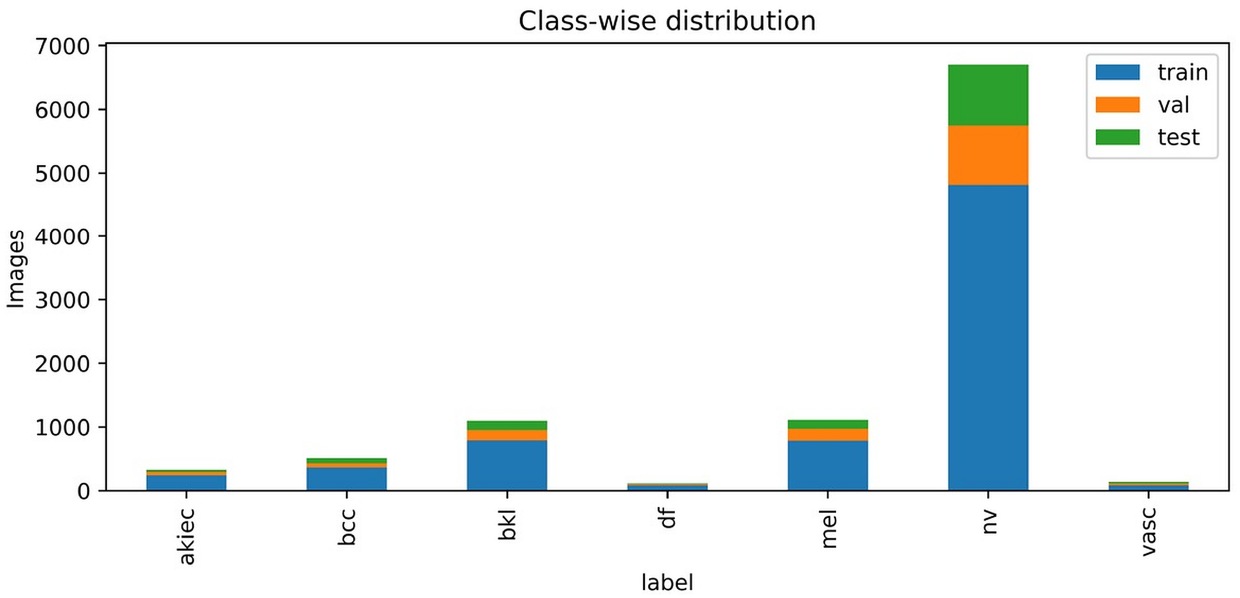
Class-wise distribution of the training, validation and test subsets.

#### External dataset

The trained model was tested, without any fine-tuning, for the robustness against domain shift with PAD-UFES-20[26] which is a collection of 2298 smartphone clinical images (after mapped labels) with patient metadata such as Fitzpatrick skin type.[35] The following classification was applied to the PAD-UFES-20 diagnoses: MEL was classified as mel, NEV as nv, BCC as bcc, ACK as akiec and SCC and SEK as bkl. There are 922 images with akiec, 845 with bcc, 244 with nv, 235 with bkl and 52 with mel in the mapped set, with no images found with df and vasc. The association of squamous cell carcinoma (SCC, including Bowen’s disease) with akiec is only approximate, so a sensitivity analysis without SCC (*n* = 2106) is also provided. To make a like-for-like comparison, the model outputs were restricted and renormalised to the five classes common to both, and the HAM10000 test set was restricted to the same classes (*n* = 1,367).

#### ISIC 2018 Task 1 (segmentation)

For the lesion segmentation experiments, ISIC 2018 Task 1 dataset[5, 6] is used, acquired from Kaggle, which consists of 2594 dermoscopic images with expert binary lesion masks. Due to the absence of an identifier for individual lesions, the pairs were randomly split (seed 43) into 1,815 training (70%), 259 validation (10%) and 520 test (20%) pairs. The segmentation network is the same for all four networks. Training and evaluation of the segmentation task is separate from the classification task: Classification networks are not fed masks, and the segmentation networks are not fed diagnostic labels.

### 3.2 Preprocessing and Augmentation

All images were transformed to RGB and resized to 224 × 224 pixels, which is the fixed size of MaxViT-Tiny’s input. The pixel values were normalised by dividing the mean and the standard deviation of the pre-trained backbone configuration. Training images were randomly resized (scale 0.7 to 1.0), horizontally and vertically flipped, rotated around a random angle (in range −20 to 20) and colour jittered (brightness 0.2, contrast 0.2, saturation 0.2, hue 0.02). Only resizing and normalising was done for validation and test images. The training subset was used to calculate class weights using the “balanced” heuristic to address the class imbalance problem, and then these weights were used in the cross entropy loss. Model selection was done with no test data; the checkpoint is the one that achieved the best validation macro-F1.

For the further leakage check, perceptual hash of each test image was compared with all training images. The minimum Hamming distance was 14 (mean 64.5), no test image was found to have a near-duplicate in the training set under the threshold of 10 bits, and test accuracy was not affected by this audit (0.869).

#### Segmentation preprocessing

Images were resized to 256 × 256 pixels (224 × 224 for Swin-T encoder which has a fixed trained size) by area-interpolation and masks were resized by nearest-neighbour interpolation and binarised at 127. Images were normalised by the mean and the standard deviation of the ImageNet data set. Training pairs were augmented by applying random horizontal and vertical flips, random 90^°^ rotations, random affine transform (rotation in ±30^°^, scale in 0.85–1.15, translation in ±8% with probability 0.5, reflective border) and brightness & contrast jitter (probability 0.8, for image only). Only size and normalisation of validation and test pairs were done.

### 3.3 Proposed Architecture

The proposed network MaxCAT-Net is a combination of a hierarchical hybrid backbone network with a light-weight attention and token-mixing head (Fig. 3, panel 3). A pretrained MaxViT-Tiny backbone *f*_*θ*_ in timm library[36] on ImageNet-1k is used to get a multi-scale feature map **F** = *f*_*θ*_ (*x*) ∈ R^512×7×7^ from a given dermoscopic image *x* ∈ R^224×224×3^. The backbone consists of a convolutional stem, and four stages, each containing 2, 2, 5 and 2 blocks with channel widths of 64, 128, 256 and 512, respectively. Each block brings together an MBConv layer for local feature extraction, and blocked (Block-SA) and grid (Grid-SA) self-attention, creating local and global receptive fields at linear complexity w.r.t the number of pixels.

**Figure 3.**
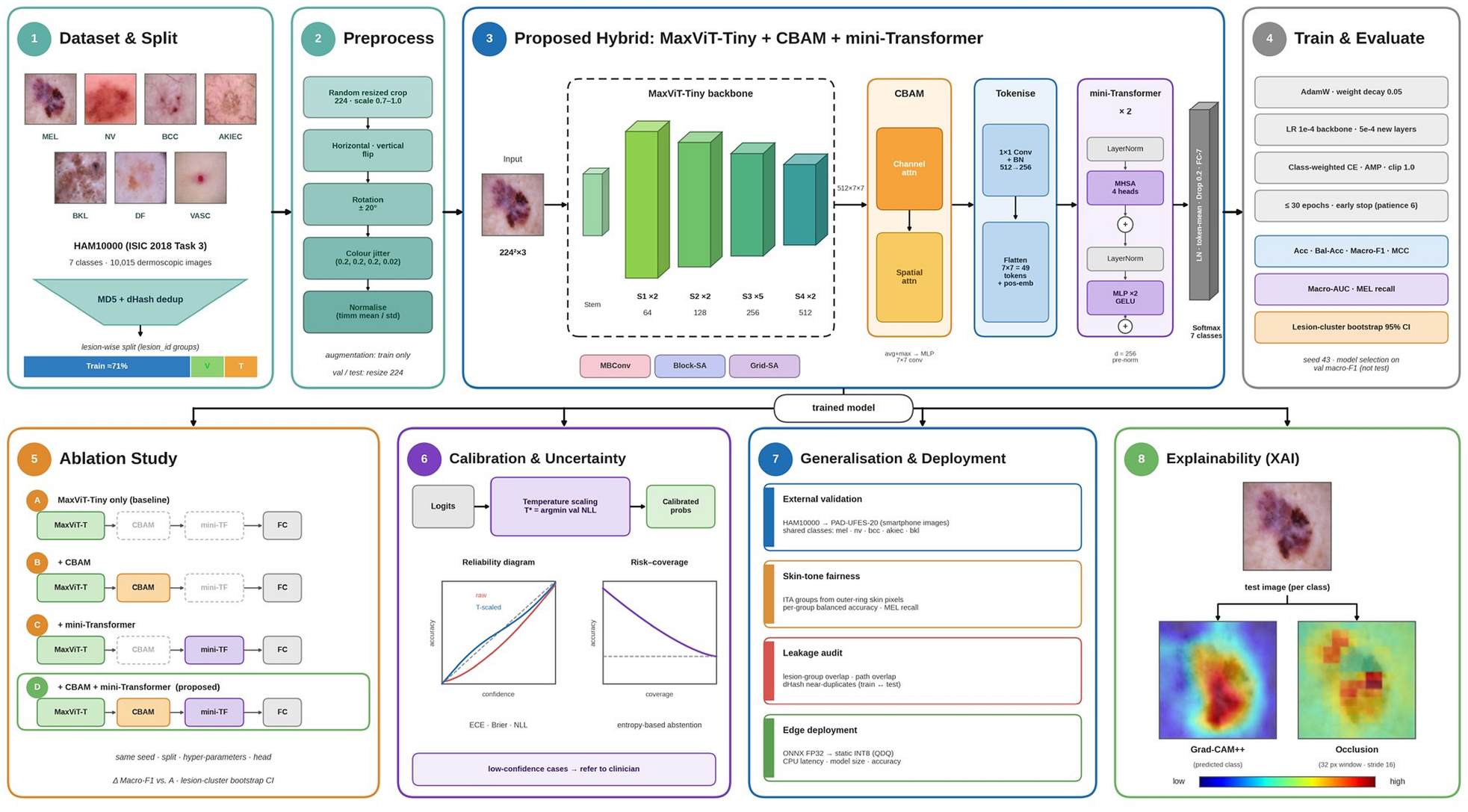
Overview of the proposed MaxCAT-Net framework: (1) dataset and lesion-wise split, (2) preprocessing, (3) MaxViT-Tiny + CBAM + mini-Transformer, (4) training and evaluation, (5) ablation study, (6) calibration and uncertainty, (7) generalisation and deployment, (8) explainability.

To emphasise lesion-relevant evidence over surrounding skin, hair and artefacts, **F** is refined by a Convolutional Block Attention Module (CBAM).[24] Channel attention computes

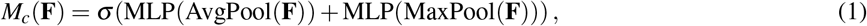

with a shared two-layer MLP of reduction ratio 16. Spatial attention computes

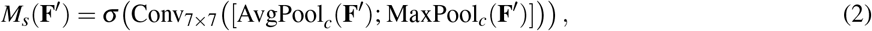

where **F**′ = *M*_*c*_(**F**) ⊙ **F**. The refined map is **F**′′ = *M*_*s*_(**F**′) ⊙ **F**′.

**F**′′ is projected to 256 channels by a 1 × 1 convolution with batch normalisation and flattened into a sequence of *N* = 49 tokens. A learnable positional embedding **E**_*pos*_ ∈ R^49×256^ is added:

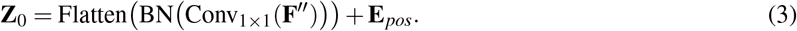

The sequence passes through *L* = 2 pre-norm Transformer encoder layers[15] with four attention heads, an MLP expansion ratio of 2 and GELU activation:

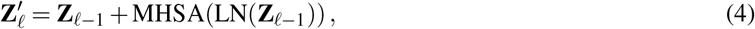

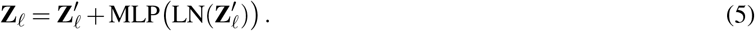

This stage re-models long-range dependencies among lesion regions after attention refinement. Finally, the tokens are layer-normalised and averaged, passed through dropout (*p* = 0.2) and a linear layer, and a softmax gives the posterior over the seven diagnostic classes:

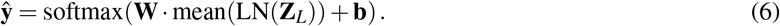

The model has 31.37 M parameters and 5.61 GMACs. All ablation variants share the same head design, seed, split and hyperparameters, so that comparisons are fair. The training objective and optimisation settings are given in Sec. 3.6.

### 3.4 Lesion Segmentation Networks

Four encoder-decoder networks were compared which output one channel logit map at the same resolution as the input. (i) Standard U-Net[9] with four down-sampling stages (feature widths 64, 128, 256, 512 and a 1,024-channel bottleneck), transposed-convolution up-sampling and skip connections, trained from random initialisation. (ii) FasterNet-T0 U-Net, which is the pretrained FasterNet-T0 encoder (with stride 4, 8, 16, 32) followed by a U-Net decoder (128/64/32/16 channels, bilinear upsampling and a prediction head). (iii) Attention U-Net[11] extends (ii) by adding an additive attention gate on each of the skip connections from the encoder, which is computed based on the feature of the decoder prior to concatenation. (iv) Swin U-Net, a combination of a pretrained Swin-T encoder[16] (swin tiny patch4 window7 224, four stages) and a convolutional U-Net decoder (widths 256, 128 and 64, followed by a 32-channel refinement block). Note that the last network is not the pure-Transformer Swin-Unet, as introduced in Ref. [14], but a Swin-encoder/U-Net-decoder hybrid. The pretrained weights for the FasterNet and Swin encoders were obtained from the ImageNet pretrained weights by timm[36] while the standard U-Net was not pretrained.

### 3.5 Evaluation Metrics

The accuracy is not useful as the test set is dominated by melanocytic nevi, so the following metrics are reported for the held out test set (*n* = 1405). Suppose TP, TN, FP and FN are true/false positives and negatives for a class.

- **Accuracy** = (TP + TN)*/*(TP + TN + FP + FN), and **balanced accuracy**, the mean per-class recall.
- **Precision, recall (sensitivity)** and **F1-score** per class, with the **macro average** (unweighted mean over classes) used as the primary summary and the weighted average reported for completeness.
- **Area under the ROC curve (AUC)** computed one-vs-rest per class and macro-averaged, plus the micro-averaged AUC.
- **Matthews correlation coefficient (MCC)**[37] for the multi-class case, which is robust to class imbalance.
- **Melanoma sensitivity** and the fraction of melanomas misclassified as nevus, as the clinically most important error.
- **Calibration:** expected calibration error (ECE), Brier score and negative log-likelihood (NLL), before and after single-parameter temperature scaling[38] (temperature fitted on the validation set).
- **Selective prediction:** accuracy on the most confident fraction (coverage) of test images.
- **Efficiency:** parameters, multiply-accumulate operations (GMACs), checkpoint size and inference latency.
- **Uncertainty:** 95% confidence intervals from a lesion-cluster bootstrap (500 resamples), in which all images of a lesion are resampled together to respect within-lesion correlation. Ablation variants share the same resamples.

#### Segmentation metrics

Per image, the predicted probabilities were thresholded at 0.5 and the resulting results compared to the reference mask using the Dice coefficient, intersection over union, precision, recall (sensitivity) and pixel accuracy which were then averaged over the 520 test images. The logits of the original, horizontally flipped and vertically flipped inputs were averaged, which is referred to as flip test-time augmentation (TTA). The non-parametric bootstrap was used to estimate the 95% confidence interval of the mean over test images (10,000 resamples). All networks were evaluated using the same set of 520 images, and thus compared with paired tests, namely: the 2-sided Wilcoxon signed-rank test with Holm correction for the 6 pairs of networks and a paired bootstrap 95% confidence interval for the mean Dice difference. The complexity of the model was reported as the number of parameters and the number of multiply-accumulate operations (GMACs) using thop with a single input and 100 forward passes averaging the GPU latencies (latencies for a batch size of 1, after 10 warm-up passes, then averaged over 100 forward passes).

### 3.6 Training Configuration

All models were trained with the same split, using the same PyTorch and timm[36] random seed (43), using deterministic cuDNN, on a single GPU (Google Colab) with mixed precision. The configuration of the proposed model and, for reference, of the baselines is given in Table 2. The proposed model adopts the differential learning rates, a lower one for pretrained backbone, and a higher one for newly initialised CBAM, Transformer and classifier layers, along with AdamW,[39] stochastic depth and gradient clipping. Training was stopped if validation macro-F1 did not improve for the patience period. The baselines were trained using a fixed learning rate and varying epochs budget (see implication in Sec. 5).

**Table 2:** Training configuration.

| Setting | MaxCAT-Net (and MaxViT ablation) | Baselines |
|---|---|---|
| Backbone | maxvit_tiny_tf_224.in1k (pretrained) | ImageNet-pretrained (timm) |
| Input size / batch size | 224 × 224 / 32 | 224 × 224 / 32 |
| Optimizer | AdamW | AdamW |
| Learning rate | 1 × 10<sup>−4</sup> (backbone), 5 × 10<sup>−4</sup> (new layers) | 3 × 10<sup>−4</sup> (all layers) |
| Weight decay | 0.05 | 1 × 10<sup>−4</sup> |
| LR scheduler | ReduceLROnPlateau (×0.5, patience 3, on val loss) | ReduceLROnPlateau (×0.5, patience 3) |
| Max epochs / early-stopping patience | 30 / 6 (val macro-F1) | 40 / 8 (val macro-F1); GhostNet hybrid 30 / 6 |
| Loss | Class-weighted cross-entropy | Class-weighted cross-entropy |
| Regularisation | Drop-path 0.1, head dropout 0.2, grad-clip 1.0 | No gradient clipping |
| Precision | Mixed precision (AMP) | Mixed precision (AMP) |
| Checkpoint selection | Best validation macro-F1 | Best validation macro-F1 |
| Seed | 43 (single run) | 43 (single run) |

#### Segmentation training

All four segmentation networks were trained with the same recipe (Table 3): AdamW[39] with a one-cycle learning-rate schedule (10% warm-up), the sum of binary cross-entropy and Dice loss, mixed precision, a maximum of 50 epochs, and early stopping with a patience of 10 epochs on validation Dice. The checkpoint with the highest validation Dice was evaluated on the test set. The networks differ in input size, batch size and (for the Swin encoder) learning rate, as listed in the table.

**Table 3:** Training configuration of the segmentation networks (ISIC 2018 Task 1, seed 43).

| **Setting** | **Standard U-Net** | **FasterNet-T0 U-Net** | **Attention U-Net** | **Swin U-Net** |
|---|---|---|---|---|
| Encoder initialisation | Random | ImageNet | ImageNet | ImageNet |
| Input size | 256 × 256 | 256 × 256 | 256 × 256 | 224 × 224 |
| Batch size | 8 | 16 | 16 | 8 |
| Peak learning rate | 3 × 10<sup>−4</sup> | 3 × 10<sup>−4</sup> | 3 × 10<sup>−4</sup> | 1 × 10<sup>−4</sup> |
| Weight decay | 1 × 10<sup>−4</sup> | 1 × 10<sup>−4</sup> | 1 × 10<sup>−4</sup> | 1 × 10<sup>−4</sup> |
| Scheduler | OneCycle | OneCycle | OneCycle | OneCycle |
| Loss | BCE + Dice | BCE + Dice | BCE + Dice | BCE + Dice |
| Max epochs / patience | 50 / 10 | 50 / 10 | 50 / 10 | 50 / 10 |
| Epochs run (selected epoch) | 50 (49) | 46 (36) | 11 (1) | 29 (19) |

## 4 Results

### 4.1 Baseline Comparison

The proposed model is compared with four baseline models on the same test split in Table 4. MaxCAT-Net achieved the highest accuracy (0.869), balanced accuracy (0.742), macro-F1 (0.751) and MCC (0.736). The single MaxViT-Tiny-CBAM baseline achieved the highest macro-AUC (0.967), while the highest sensitivity to melanoma was achieved by MobileNetV3-CBAM (0.601), which was the same as single MaxViT baseline. Baselines CNN and GhostNet hybrid were 3.6–6.3 accuracy points behind the proposed model. The difference between the macro-F1 confidence intervals is not statistically significant (e.g. proposed 0.691–0.799 and ResNet152-CBAM 0.589–0.712).

**Table 4:** Baseline comparison on the HAM10000 test set (*n* = 1,405; seed 43). Best performance value per metric column in bold.

| **Model** | **Acc.** | **Bal. Acc.** | **Macro-F1** | **AUC** | **MEL recall** | **Params (M)** | **GMACs** | **Size (MB)** |
|---|---|---|---|---|---|---|---|---|
| ResNet152-CBAM | 0.821 | 0.654 | 0.654 | 0.957 | 0.594 | 58.68 | 11.60 | 235.7 |
| MobileNetV3-L-CBAM | 0.833 | 0.652 | 0.660 | 0.952 | **0.601** | 4.33 | 0.22 | 17.5 |
| GhostNet-100 + CBAM + Transformer | 0.806 | 0.651 | 0.650 | 0.942 | 0.566 | 3.18 | 0.17 | 13.0 |
| MaxViT-Tiny-CBAM (single) | 0.851 | 0.710 | 0.728 | **0.967** | **0.601** | 30.44 | 5.33 | 122.2 |
| MaxCAT-Net (proposed) | **0.869** | **0.742** | **0.751** | 0.944 | 0.587 | 31.37 | 5.61 | 125.9 |

Latency was measured in separate sessions on possibly different GPUs and is therefore not compared across models; for the proposed model it was 51.9 ms per image (GPU) and 382.5 ms (single CPU thread).

### 4.2 Ablation Study

Figure 4 and Table 5 show the ablation of the two extra components on the MaxViT-Tiny backbone under the same conditions. Adding CBAM, the Transformer encoder, or both improved test accuracy (0.856 → 0.864–0.869), balanced accuracy (0.703 → 0.724–0.742) and macro-F1 (0.734 → 0.751–0.756) relative to the plain backbone. The best macro-F1 was obtained by MaxViT+CBAM (0.756), whereas MaxCAT-Net had the highest accuracy and balanced accuracy and the highest validation macro-F1 was obtained by MaxViT+Transformer (0.802). These gains should be viewed as a trend as all F1 confidence intervals overlap, therefore these gains cannot be considered a statistically significant effect. The proposed variant was MaxCAT-Net (full hybrid) which was chosen a priori. The simpler variants achieved the highest Macro-AUC (Macro-AUC of 0.972 for MaxViT+CBAM) and melanoma recall (Melanoma Rec 0.643 for the plain backbone).

**Table 5:** Ablation of CBAM and the Transformer encoder on MaxViT-Tiny (test set).

| **Variant** | **CBAM** | **Trans.** | **Params (M)** | **Val F1** | **Acc.** | **Bal. Acc.** | **Macro-F1** | **F1 95% CI** | **AUC** | **MEL rec.** |
|---|---|---|---|---|---|---|---|---|---|---|
| MaxViT only | – | – | 30.14 | 0.783 | 0.856 | 0.703 | 0.734 | 0.668–0.788 | 0.957 | 0.643 |
| + CBAM | ✓ | – | 30.18 | 0.770 | 0.864 | 0.724 | 0.756 | 0.690–0.808 | 0.972 | 0.601 |
| + Transformer | – | ✓ | 31.34 | 0.802 | 0.868 | 0.726 | 0.753 | 0.690–0.802 | 0.947 | 0.552 |
| + CBAM + Transformer (MaxCAT-Net) | ✓ | ✓ | 31.37 | 0.788 | 0.869 | 0.742 | 0.751 | 0.690–0.801 | 0.944 | 0.587 |

**Figure 4.**
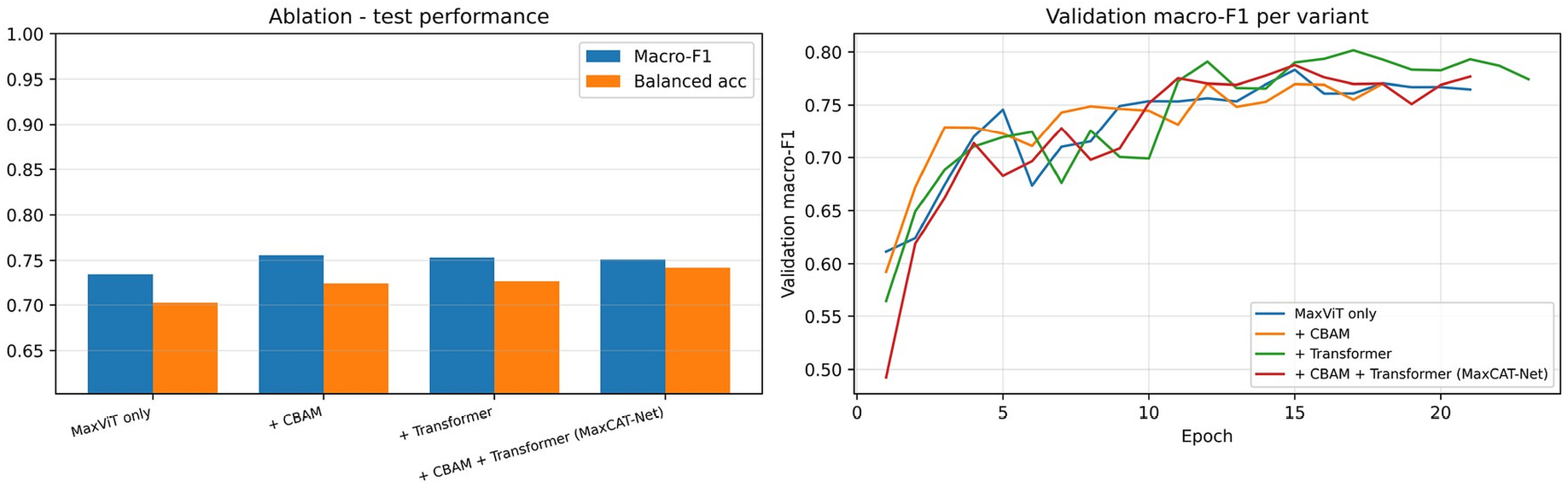
Ablation results: test macro-F1 and balanced accuracy (left) and validation macro-F1 per epoch for each variant (right).

### 4.3 Class-wise Performance and ROC Analysis

The class-wise results are given in Table 6. Melanocytic nevi were recognised with the highest F1 (0.937) and recall (0.956). Basal cell carcinoma (F1 0.789), dermatofibroma (0.783) and vascular lesions (0.769) were recognised reliably despite their small support, while actinic keratoses had the lowest F1 (0.588) and melanoma had a recall of 0.587 with F1 of 0.646. Per-class AUC values lay between 0.925 and 0.970, and the micro-averaged AUC was 0.978 (Fig. 5). The overall Cohen’s kappa was 0.735.

**Table 6:** Class-wise performance of the proposed model on the test set.

| **Class** | **Precision** | **Recall** | **F1-score** | **Support** | **AUC** |
|---|---|---|---|---|---|
| akiec | 0.606 | 0.571 | 0.588 | 35 | 0.946 |
| bcc | 0.767 | 0.812 | 0.789 | 85 | 0.969 |
| bkl | 0.788 | 0.708 | 0.746 | 147 | 0.925 |
| df | 0.750 | 0.818 | 0.783 | 11 | 0.921 |
| mel | 0.718 | 0.587 | 0.646 | 143 | 0.945 |
| nv | 0.919 | 0.956 | 0.937 | 957 | 0.970 |
| vasc | 0.800 | 0.741 | 0.769 | 27 | 0.929 |
| **Macro average** | **0.764** | **0.742** | **0.751** | **1,405** | **0.944** |
| Weighted average | 0.864 | 0.869 | 0.865 | 1,405 | – |

**Figure 5.**
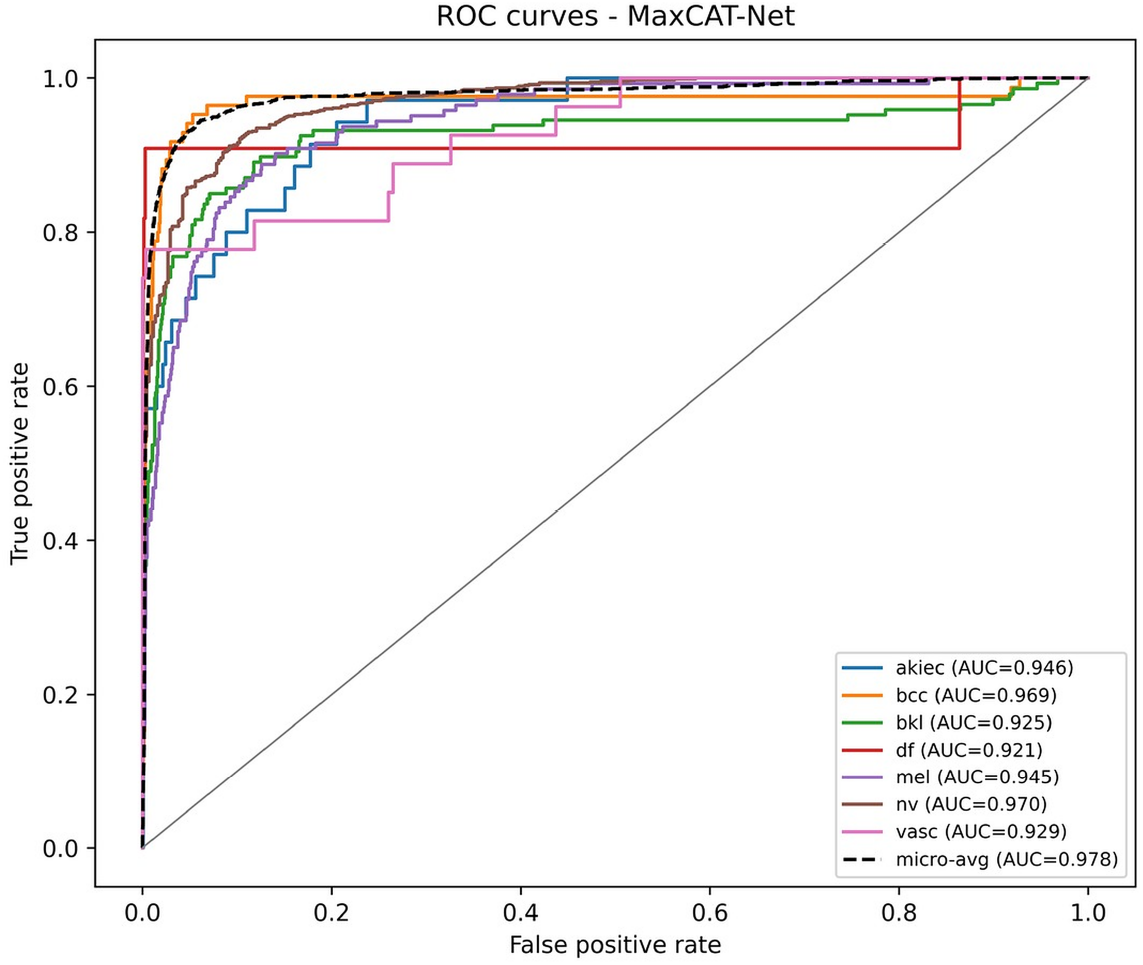
One-vs-rest ROC curves of the proposed model on the test set. malignant and 915 of 957 nevi were correctly classified.

## 4.4 Confusion Matrix

The confusion matrix is shown in Fig. 6. Of the 143 melanomas, 84 were correctly classified while 51 (35.7%) were classified as nevus, the most common error and clinical concern. Other common errors included akiec predicted as bkl (8 of 35) and bkl predicted as nv (19 of 147) or mel (13 of 147). On the other hand, 17 nevi were also misclassified as malignant and 915 of 957 nevi were correctly classified.

**Figure 6.**
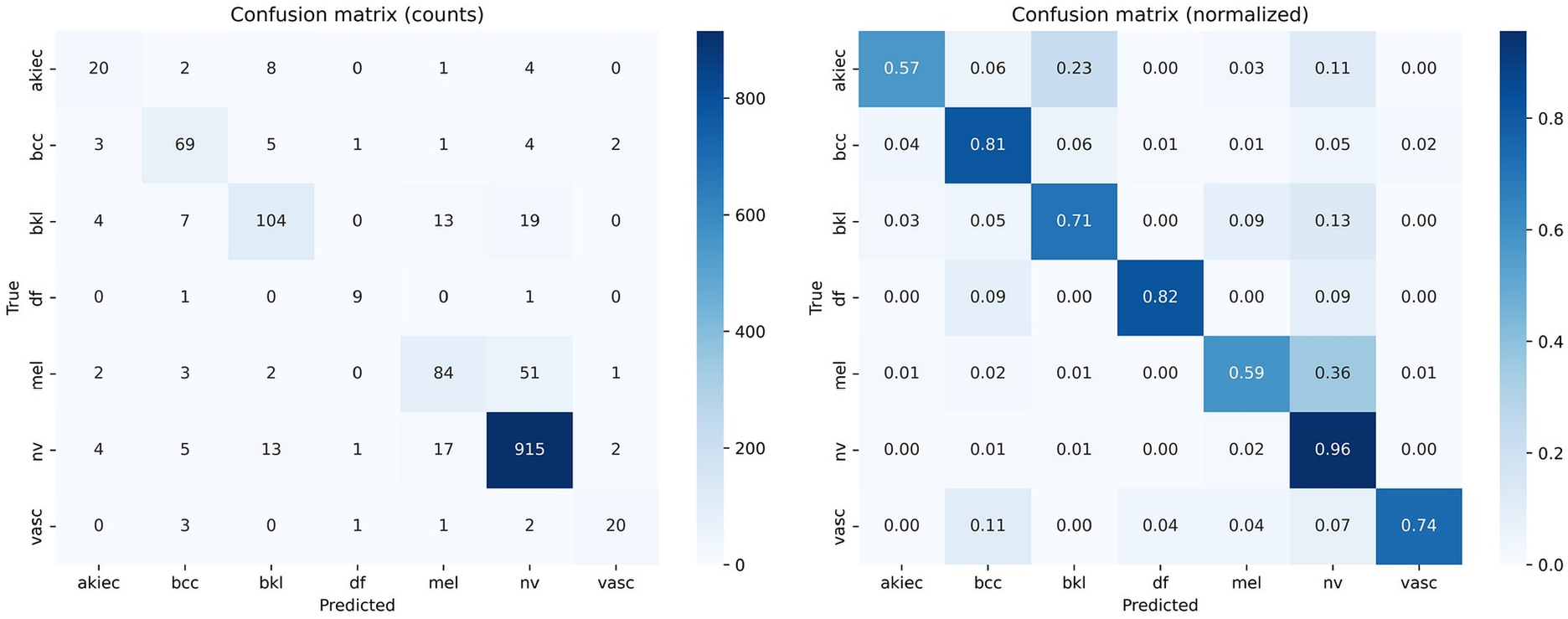
Confusion matrix of MaxCAT-Net on the test set: counts (left) and row-normalised (right).

### 4.5 Training and Validation Curves

The results of training and validation accuracy as well as loss and validation macro-F1 is depicted in Fig. 7. It was stopped after 6 epochs with no improvement and the best checkpoint was used (epoch 15, validation macro-F1 0.788).

**Figure 7.**
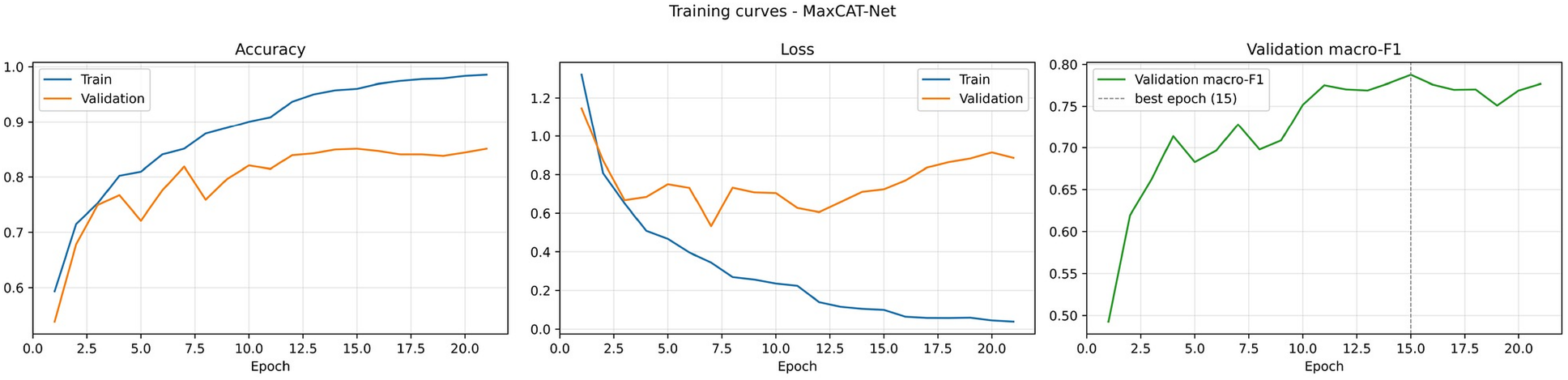
Training and validation accuracy (left), loss (centre) and validation macro-F1 (right); the dashed line marks the selected epoch.

Training accuracy kept increasing to 0.986 while training loss decreased to 0.039, while validation accuracy remained at around 0.84–0.85, and validation loss after the minimum at epoch 7 (0.533) increased to 0.886 at the last epoch. This increasing disparity shows that the later epochs have started to over-fit on the minority classes which is corrected by early stopping on validation macro-F1

### 4.6 Calibration and Selective Prediction

The class-weighted training lead to overconfident predictions (ECE 0.099). A single temperature (*T* = 2.067), fitted on the validation set, reduced ECE to 0.038, the Brier score from 0.224 to 0.200 and the NLL from 0.661 to 0.444 (Table 7, Fig. 8). In Table 7 and Fig. 9, the accuracy of the predictions on the cases not deferred is monotonically increasing from 0.869 at full coverage to 0.915, 0.945 and 0.959 at 90%, 80% and 70% coverage, respectively, although the increase is very slight after 50% coverage (0.966). This indicates that confidence may be a tool for identifying cases for specialist review.

**Table 7:** Calibration metrics (left) and selective prediction (right).

| **Metric** | **Raw** | **Temp. scaled (*T* = 2.067)** | **Coverage** | **Accuracy on covered** |
|---|---|---|---|---|
| ECE | 0.0994 | 0.0379 | 1.0 | 0.869 |
| Brier score | 0.2239 | 0.1996 | 0.9 | 0.915 |
| NLL | 0.6613 | 0.4443 | 0.8 | 0.945 |
|  |  |  | 0.7 | 0.959 |
|  |  |  | 0.5 | 0.966 |

**Figure 8.**
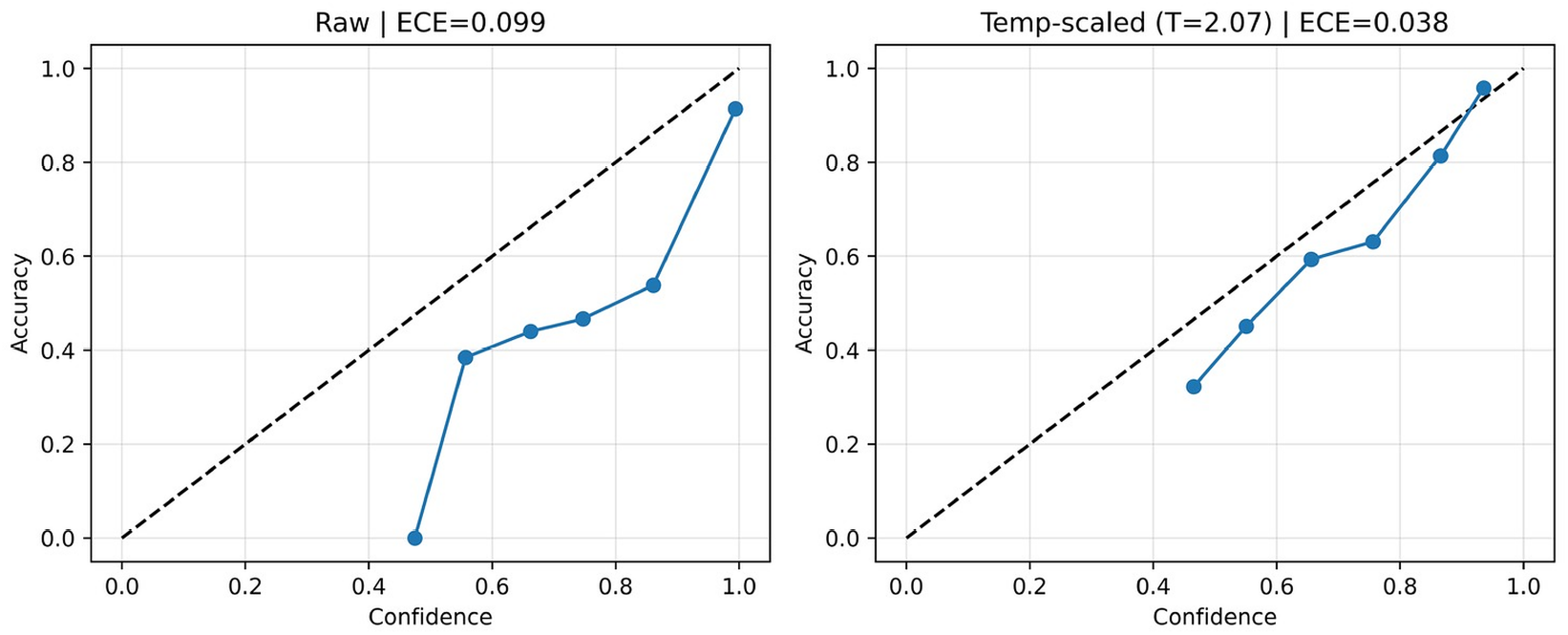
Reliability diagrams before and after temperature scaling.

**Figure 9.**
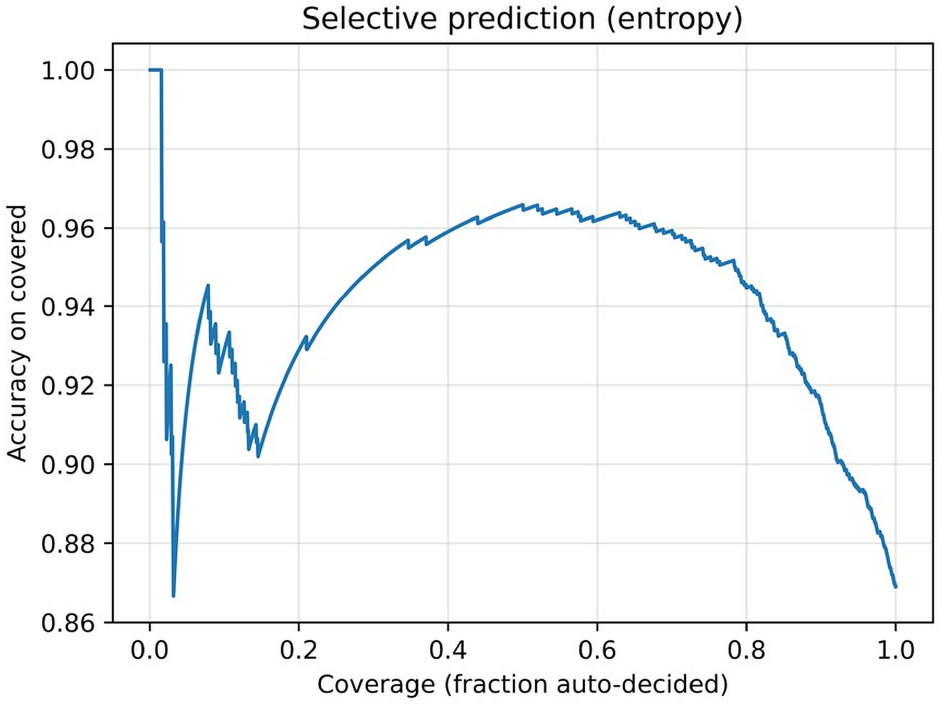
Risk–coverage (accuracy versus coverage) curve for selective prediction.

### 4.7 External Validation on PAD-UFES-20

The results on the external smartphone-image dataset dropped significantly (Table 8). On the five shared classes, accuracy fell from 0.875 (internal) to 0.341 (external), balanced accuracy from 0.733 to 0.353, macro-F1 from 0.746 to 0.253 and macro-AUC from 0.952 to 0.680. On the external data, the model proved to be inaccurate and overly confident, with a lower score for melanoma recall (0.594 to 0.096) and a higher score for ECE (0.094 to 0.569). However, the accuracy of the akiec mapping was not significantly affected when the SCC was removed from the mapping (accuracy 0.360, balanced accuracy 0.349), suggesting that the effect of the label-mapping approximation is not the primary reason for the loss in accuracy. The external confusion matrix is displayed in Fig. 10.

**Table 8:** Internal (HAM10000 test, shared classes) versus external (PAD-UFES-20) performance of the proposed model.

| **Evaluation set** | *n* | **Acc.** | **Bal. Acc.** | **Macro-F1** | **AUC** | **MEL recall** | **ECE** |
|---|---|---|---|---|---|---|---|
| HAM10000 test (internal) | 1,367 | 0.875 | 0.733 | 0.746 | 0.952 | 0.594 | 0.094 |
| PAD-UFES-20 (external) | 2,298 | 0.341 | 0.353 | 0.253 | 0.680 | 0.096 | 0.569 |
| PAD-UFES-20 (SCC excluded) | 2,106 | 0.360 | 0.349 | 0.258 | 0.679 | 0.096 | 0.552 |
| Drop (internal − external) | – | 0.534 | 0.381 | 0.494 | 0.273 | 0.498 | –0.474 |

**Figure 10:**
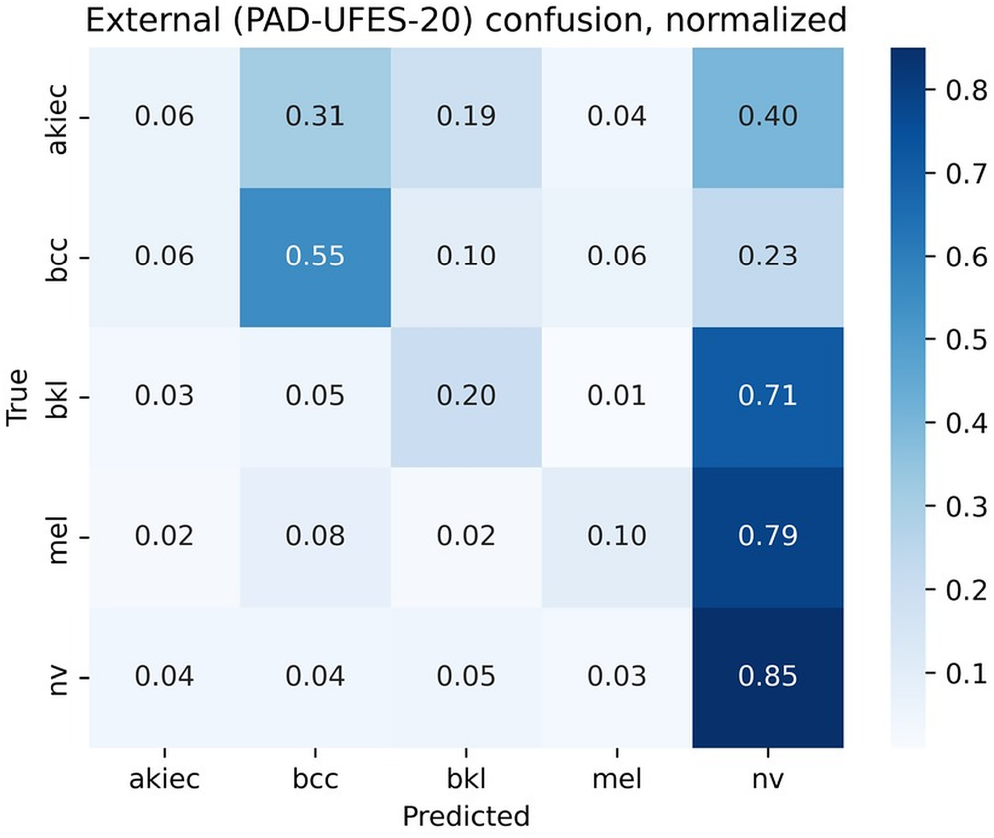
Confusion matrix of the proposed model on PAD-UFES-20 (shared classes).

The performance of the proposed model was compared with the other trained models in the same external set as shown in Table 9. The external accuracy of all the models were in the range of 0.263 to 0.366 and melanoma recall ranged from 0.077 to 0.308. ResNet152-CBAM baseline showed the highest external accuracy (0.366) and balanced accuracy (0.369), MaxCAT-Net showed the highest external AUC (0.680), and the GhostNet hybrid had the highest external melanoma recall (0.308) corresponding to the lowest accuracy. Hence, no model can be claimed to be robust based on this experiment. An external evaluation has not been performed for MobileNetV3-CBAM.

**Table 9:** External validation (PAD-UFES-20, *n* = 2,298) of the trained models.

| **Model** | **Acc.** | **Bal. Acc.** | **Macro-F1** | **AUC** | **MEL recall** | **ECE** |
|---|---|---|---|---|---|---|
| ResNet152-CBAM | 0.366 | 0.369 | 0.297 | 0.666 | 0.077 | 0.389 |
| GhostNet-100 + CBAM + Transformer | 0.263 | 0.342 | 0.215 | 0.646 | 0.308 | 0.588 |
| MaxViT-Tiny-CBAM (single) | 0.316 | 0.346 | 0.240 | 0.640 | 0.173 | 0.553 |
| MaxCAT-Net (proposed) | 0.341 | 0.353 | 0.253 | 0.680 | 0.096 | 0.569 |

#### Subgroup analysis

Performance by skin-tone-related groups is reported in Table 10. The individual typology angle (ITA) calculated from the image as a proxy for skin tone was able to be estimated in 64.9% of the test images, with the worst to best balanced-accuracy difference between groups with at least 30 images being 0.176 (very light 0.712 versus tan 0.889). Few images are present in the medium tone groups, and there are very few melanomas (*n* = 135, 84 and 31), thus giving wide confidence intervals, and the brown group (*n* = 10) is too small for interpretation. With PAD-UFES-20, the accuracy ranged from 0.387 to 0.425 for Fitzpatrick type (FT) I–IV, and from 0.318 to 0.411 for balanced accuracy with no group performing well. Fitzpatrick types V and VI (*n* = 10 and 1) were too low in numbers to analyze.

**Table 10:** Subgroup performance. Top: HAM10000 test by ITA group. Bottom: PAD-UFES-20 by Fitzpatrick type.

| **Group** | *n* | **Accuracy** | **Balanced acc.** | **Macro-F1** |
|---|---|---|---|---|
| ITA very light | 652 | 0.863 | 0.712 | 0.720 |
| ITA light | 135 | 0.970 | 0.869 | 0.851 |
| ITA intermediate | 84 | 0.917 | 0.717 | 0.705 |
| ITA tan | 31 | 0.968 | 0.889 | 0.822 |
| ITA brown | 10 | 0.800 | 0.667 | 0.625 |
| Fitzpatrick I | 153 | 0.425 | 0.327 | 0.194 |
| Fitzpatrick II | 876 | 0.390 | 0.411 | 0.240 |
| Fitzpatrick III | 392 | 0.416 | 0.380 | 0.255 |
| Fitzpatrick IV | 62 | 0.387 | 0.318 | 0.236 |

### 4.8 Explainability and Deployment Profile

For the test images with the highest confidence for each class, gradient-weighted class activation maps (Grad-CAM++)[40] and occlusion sensitivity maps were created (Fig. 11). The activation for melanoma had its focus on the lesion itself and/or one or more typical lesion structures, whereas the most activated portion of the image for nevus and dermatofibroma primarily occurred outside the lesion body, close to the border of the image or a small adjacent area, indicating that the model may be relying on non-lesion cues for this class. An explanation with a SHAP object was tried but failed due to GPU memory problems, and is not reported.

**Figure 11:**
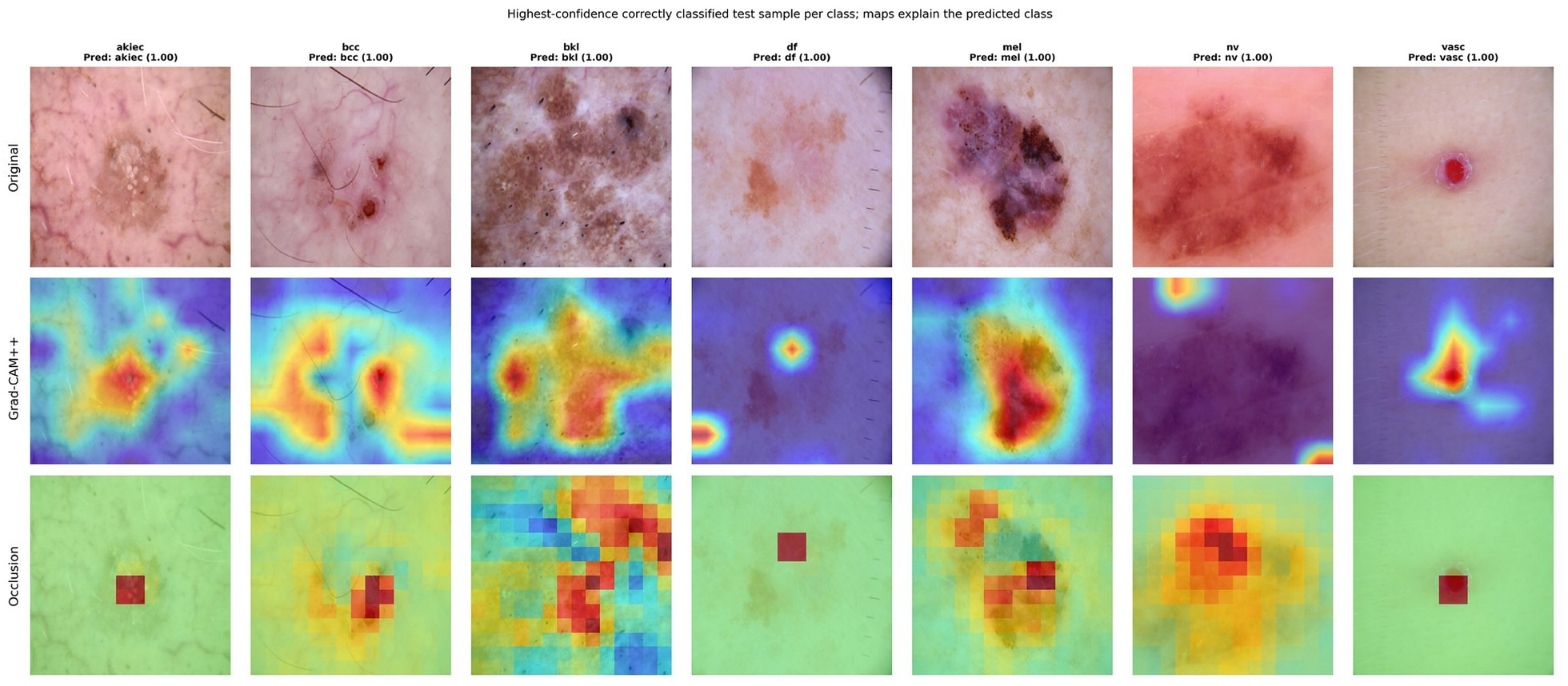
Original images, Grad-CAM++ and occlusion maps for one correctly classified test sample per class. procedure. FP16 weights (62.7 MB) were not evaluated for accuracy.

Table 11 summarises the deployment profile. Converting the model to ONNX (FP32) preserved accuracy exactly (0.869) and gave a CPU latency of 242 ms per image under ONNX Runtime, compared with 383 ms for single-thread PyTorch. Static INT8 post-training quantisation shrank the file from 126 MB to 36 MB but caused a collapse in performance (accuracy 0.483, macro-F1 0.194), so quantised deployment is not supported by the current calibration

**Table 11:** Deployment profile of the proposed model.

| **Format** | **Size (MB)** | **CPU latency (ms)** | **Test accuracy** | **Test macro-F1** |
|---|---|---|---|---|
| PyTorch FP32 checkpoint | 125.9 | 382.5 (1 thread) | 0.869 | 0.751 |
| ONNX FP32 | 126.5 | 242.1 | 0.869 | 0.751 |
| ONNX INT8 (static) | 35.8 | 242.6 | 0.483 | 0.194 |

### 4.9 Lesion Segmentation on ISIC 2018 Task 1

The results of the four networks in terms of segmentation accuracy for the 520 held-out test images are presented in Table 12. The Swin U-Net obtained the highest Dice (0.903; 95% CI 0.893–0.912) and IoU (0.838), followed closely by the FasterNet-T0 U-Net (Dice 0.901), the Attention U-Net (0.900) and, at a clear distance, the standard U-Net (0.888). There are at most 0.003 difference in Dice between the three pretrained-encoder networks, and they have overlapping confidence intervals, while the standard U-Net is 0.012 to 0.015 lower. All models showed pixel accuracy of 0.956–0.963, but this was due to the vast area of the background. Precision (0.918–0.922) was higher than recall (0.891–0.911) in all networks, and errors were slightly more likely to be under-than over-segmentations. For all networks, Flip-TTA was able to boost Dice by 0.0015 to 0.0024 without altering the network ranking.

**Table 12:** Segmentation results on the ISIC 2018 Task 1 test set (*n* = 520; mean per-image values, seed 43). Best value per column in bold.

| Model | Dice (95% CI) | IoU | Precision | Recall | Pixel acc. | Dice, flip-TTA | IoU, flip-TTA |
|---|---|---|---|---|---|---|---|
| Standard U-Net | 0.888 (0.877–0.899) | 0.817 | 0.920 | 0.891 | 0.956 | 0.890 | 0.818 |
| FasterNet-T0 U-Net | 0.901 (0.890–0.910) | 0.834 | 0.922 | 0.907 | 0.962 | 0.903 | 0.837 |
| Attention U-Net (FasterNet-T0) | 0.900 (0.889–0.910) | 0.834 | 0.918 | 0.911 | 0.962 | 0.902 | 0.836 |
| **Swin U-Net** | **0.903 (0.893–0.912)** | **0.838** | **0.922** | **0.911** | **0.963** | **0.905** | **0.840** |

Table 13 summarises computational cost and training behaviour. The FasterNet-T0 U-Net and the Attention U-Net have about 3.1–3.2 M parameters, a 12.6–12.8 MB checkpoint and below 1 GMAC, which is roughly ten times fewer parameters than the Swin U-Net and the standard U-Net and about 57 times fewer multiply-accumulate operations than the standard U-Net (54.7 GMACs). The FasterNet-T0 U-Net also had the lowest GPU latency (5.3 ms), and the standard U-Net the highest (43.8 ms). The GMACs of the Swin U-Net were not recorded. Compared with the Swin U-Net, the FasterNet-T0 U-Net gave up 0.0024 Dice for a ten-fold reduction in parameters.

**Table 13:** Complexity and training behaviour of the segmentation networks.

| **Model** | **Params (M)** | **GMACs** | **GPU latency (ms)** | **Size (MB)** | **Epochs run** | **Selected epoch** |
|---|---|---|---|---|---|---|
| Standard U-Net | 31.04 | 54.74 | 43.8 | 124.2 | 50 | 49 |
| FasterNet-T0 U-Net | 3.12 | 0.96 | 5.3 | 12.6 | 46 | 36 |
| Attention U-Net (FasterNet-T0) | 3.17 | 0.99 | 6.4 | 12.8 | 11 | 1 |
| Swin U-Net | 31.62 | n/a | 13.4 | 126.6 | 29 | 19 |

Latency was measured in separate Colab sessions and with different input sizes (224 × 224 for Swin U-Net, 256 × 256 otherwise), so it is indicative only. GMACs for the Swin U-Net were not computed.

### 4.10 Statistical Comparison of Segmentation Networks

As the four networks were tested on the same set of test images, paired tests on per-image dice were possible (Table 14). Each of the three pretrained-encoder networks was significantly superior to the standard U-Net, with mean Dice differences ranging from −0.012 to −0.015, and paired bootstrap 95% confidence intervals not containing zero, and Holm-adjusted Wilcoxon *p*-values less than 0.002. No statistically significant difference was detected between the Attention U-Net and the FasterNet-T0 U-Net (ΔDice = −0.0007, 95% CI −0.0033 to +0.0017, adjusted *p* = 0.92), suggesting that the attention gates were not measurably beneficial. The two tests produced conflicting results: the Wilcoxon test which compares the distribution of paired differences was significant (adjusted *p* = 0.007 against each); the bootstrap confidence intervals for the difference in the mean Dice (−0.0038 to +0.0085 against FasterNet-T0 U-Net) contained zero. The Swin U-Net thus has a slight advantage with a typical image, but the difference in mean Dice, driven by a few failures, is statistically indistinguishable from 0 and is not clinically significant (0.002–0.003).

**Table 14:** Paired comparison of per-image Dice between segmentation networks (*n* = 520). Δ = mean Dice of the first minus the second model; *p*-values are from the two-sided Wilcoxon signed-rank test.

| **Comparison (A vs B)** | **Δ Dice** | **95% CI of Δ** | *p* **(Wilcoxon)** | *p* **(Holm)** |
|---|---|---|---|---|
| Swin U-Net vs Standard U-Net | +0.0149 | +0.0079 to +0.0225 | 2.3 × 10<sup>−10</sup> | 1.4 × 10<sup>−9</sup> |
| FasterNet-T0 U-Net vs Standard U-Net | +0.0125 | +0.0052 to +0.0203 | 3.6 × 10<sup>−4</sup> | 1.4 × 10<sup>−3</sup> |
| Attention U-Net vs Standard U-Net | +0.0118 | +0.0042 to +0.0198 | 1.5 × 10<sup>−4</sup> | 7.3 × 10<sup>−4</sup> |
| Swin U-Net vs FasterNet-T0 U-Net | +0.0024 | −0.0038 to +0.0085 | 2.9 × 10<sup>−3</sup> | 7.4 × 10<sup>−3</sup> |
| Swin U-Net vs Attention U-Net | +0.0031 | −0.0028 to +0.0090 | 2.5 × 10<sup>−3</sup> | 7.4 × 10<sup>−3</sup> |
| FasterNet-T0 U-Net vs Attention U-Net | +0.0007 | −0.0017 to +0.0033 | 0.92 | 0.92 |

### 4.11 Training Curves and Qualitative Results

The training and validation behaviour is displayed in Fig. 12. The standard U-Net trained from random initialisation converged slowly and was not yet fully converged at the 50-epoch limit with a validation Dice of 0.880 at the selected epoch, which may thus be under-estimating the performance of a longer training. Both the Swin U-Net and the FasterNet-T0 U-Net achieved a validation Dice score of approximately 0.89 following 15–20 epochs of training, but failed to continue improving beyond this point. The logged training history for the Attention U-Net is quite distinct from the others: the training loss at the first logged epoch (0.16) was already quite close to the end of the training loss of the other networks (0.16–0.24), the best validation Dice was found at the first logged epoch, and early stopping had ended the training run after 11 epochs. This indicates that the logged run originated from weights which already had been trained (e.g., the training cell was run the second time in the same session), and the history of this run is not comparable to the others (see Sec. 5).

**Figure 12:**
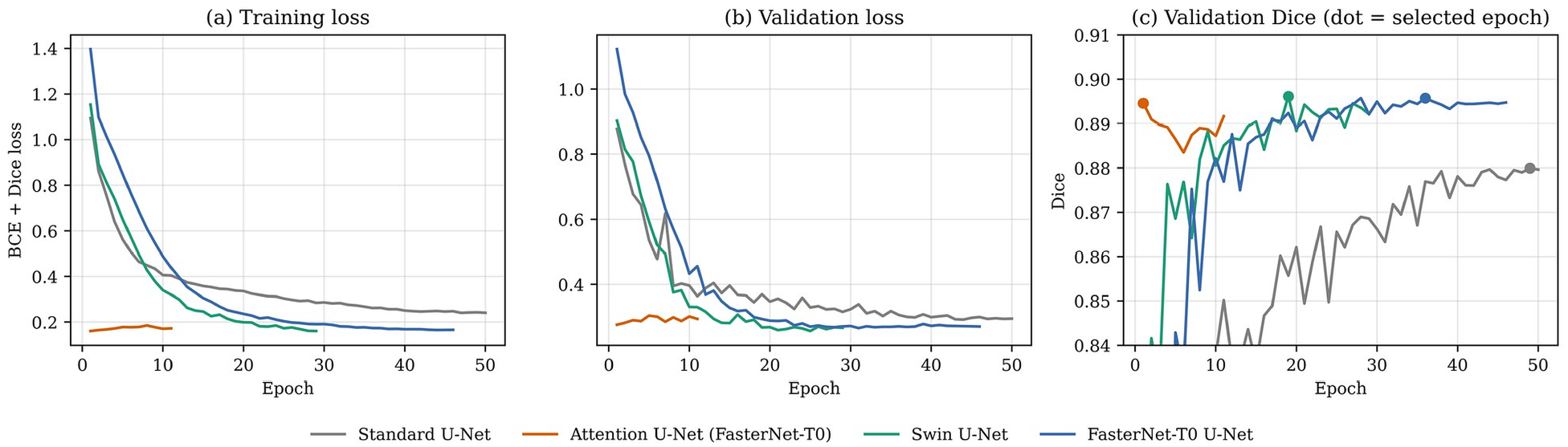
Training loss, validation loss and validation Dice of the four segmentation networks; dots mark the selected epoch.

Per-image Dice was highly skewed: the median Dice was 0.929–0.938 for all networks, whereas 12–15 test images (2.3–2.9%) had Dice below 0.5 and 42–63 images (8–12%) had Dice below 0.8. Five of these images achieved Dice below 0.5 for all four networks, and the per-image Dice values of the networks were well correlated (Pearson *r* = 0.73–0.97), thus most failures are common and likely due to the images or their masks and are not necessarily caused by a specific architecture. The FasterNet-T0 U-Net on the 2 lowest scoring test images, a median and the highest-scoring image is shown in Fig. 13. The two failures where the reference mask is atypical (ring-shaped annotation around the lesion) and the lesion is faint and low-contrast with a ruler in view and the prediction covers only part of the annotated region, thus indicating that annotation inconsistency and low contrast dominate the error, rather than boundary quality. Median and best cases are close together on the lesion border.

**Figure 13:**
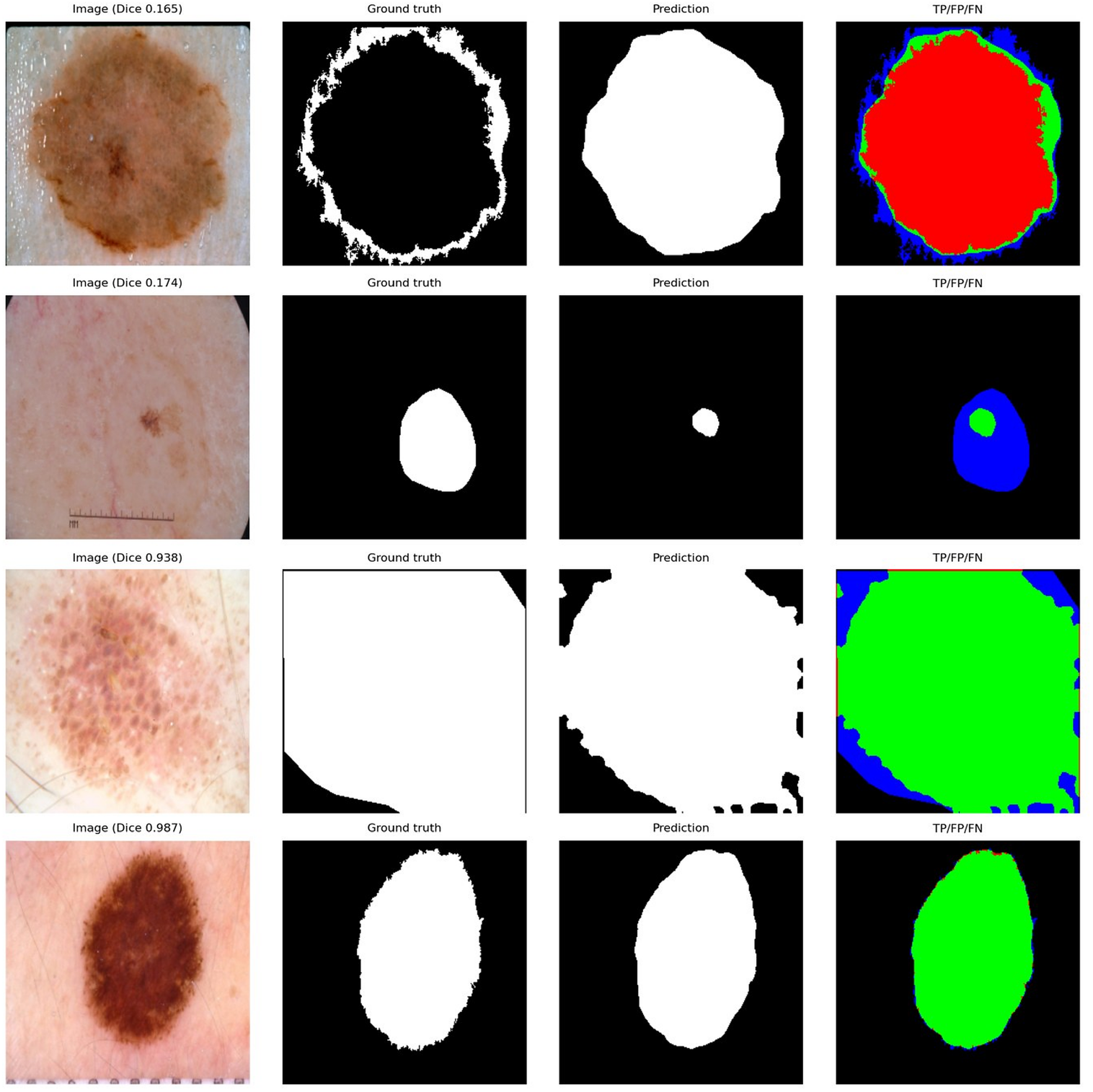
Qualitative segmentation results of the FasterNet-T0 U-Net: image, reference mask, prediction and overlay (green = true positive, red = false positive, blue = false negative), for two of the lowest-scoring cases (Dice 0.165 and 0.174), a median case (0.938) and the highest-scoring case (0.987).

## 5 Discussion

### 5.1 Main Findings

The best results in terms of accuracy, balanced accuracy, macro-F1 and MCC were obtained by the hybrid model MaxCAT-Net (MaxViT-Tiny + CBAM + Transformer) on a leakage-controlled split. This represents a 3.6–6.3 point increase in accuracy compared to CNN baselines (0.821–0.833) and the lightweight GhostNet hybrid (0.806), and a significantly larger increase in balanced accuracy (0.742 versus 0.652–0.654), reflecting that the improvement is primarily driven by the recognition of minority classes rather than the dominant nevus class. These numbers are probably lower, but more correct, than numbers obtained via image-level random splits due to leakage controls (deduplication, lesion-wise splitting and perceptual-hash audit).

### 5.2 Impact of the Additional Modules

All the variants outperformed the plain MaxViT-Tiny backbone, but the macro-F1 improvements of +0.017 to +0.022 are rather small considering the width of the confidence intervals, and the macro-F1 of the simpler MaxViT+CBAM (0.756) is similar to MaxCAT-Net (0.751), while being superior in terms of AUC and the number of parameters (just 0.2M less). The gain of MaxCAT-Net is thus limited to accuracy and balanced accuracy and should be verified with several seeds. Secondly, the advantage of CBAM is relative to the backbone. In the GhostNet ablation (Table 15), the test accuracy (0.813 → 0.779) and macro-F1 (0.657 → 0.636) were improved when the CNN was not used. The best GhostNet model was when the CNN was combined with Transformer (accuracy 0.832, macro-F1 0.687). The full GhostNet hybrid (0.806 / 0.650) was not better than the plain CNN on macro-F1. The value of attention modules thus has to be tested on every backbone and does not help uniformly.

**Table 15:** Ablation on the GhostNet-100 backbone (test set, seed 43).

| **Variant** | **CBAM** | **Trans.** | **Acc.** | **Bal. Acc.** | **Macro-F1** | **F1 95% CI** | **AUC** |
|---|---|---|---|---|---|---|---|
| CNN only | – | – | 0.813 | 0.662 | 0.657 | 0.586–0.716 | 0.949 |
| + CBAM | ✓ | – | 0.779 | 0.667 | 0.636 | 0.572–0.690 | 0.945 |
| + Transformer | – | ✓ | 0.832 | 0.686 | 0.687 | 0.621–0.742 | 0.941 |
| + CBAM + Transformer | ✓ | ✓ | 0.806 | 0.651 | 0.650 | 0.585–0.702 | 0.942 |

### 5.3 Comparisons with Baselines and a Caveat

The baselines and the proposed model were trained with varying hyperparameters (Table 2): the MaxViT backbone has a lower backbone learning rate, and it has stronger weight decay, drop-path and gradient clipping, while the baselines have a single learning rate. However, the improvement of MaxCAT-Net compared with the single MaxViT-Tiny-CBAM baseline (accuracy 0.869 vs. 0.851, macro-F1 0.751 vs. 0.728) is composed of the architectural change and different training recipe. The controlled evidence for the architecture is the ablation (Table 5), which yielded a plain backbone under the same recipe with 0.856 accuracy and 0.734 macro-F1. This should be borne in mind when considering the baseline comparison.

### 5.4 Melanoma Sensitivity

Although the overall scores were the highest, melanoma recall was 0.587 and 35.7% of melanomas were classified as nevi. It is the greatest limitation of a screening application, which all models share (recall 0.566–0.601). The higher accuracy of the hybrid is in fact partly due to the majority class as the rates of melanoma recall were highest for the plain backbone (0.643) and lowest for the Transformer variant (0.552). It may be worth looking at class-weighted cross-entropy, focal loss, decision threshold tuning or cost sensitive objectives to sacrifice some specificity for sensitivity.

### 5.5 Calibration and Triage

Temperature scaling led to a reduction of 62% in ECE while maintaining the same predictions, and selective prediction resulted in a 94.5% accuracy at 80% coverage, using the Brier score and NLL. These properties enable a triage system where patients’ cases of lower confidence may be referred to a dermatologist. Class weights are used in training which cause the class prior to change; therefore, there is no single global temperature that can correct the calibration of every class.

### 5.6 Generalisation and Fairness

The external validation is the most sobering result. A model with 0.875 internal accuracy on the shared classes scored only 0.341 on PAD-UFES-20 and all other models scored similarly (0.263–0.366). There are several reasons for this drop, such as changes in illumination, resolution, field of view and background (from dermoscopy to smartphone photography), differences in the case mix (PAD-UFES-20 has many more bcc and akiec lesions, and few nevi) and the approximate label mapping (no df or vasc classes). The sensitivity analysis without SCC indicates that the mapping is not the major contributor. In addition, the subgroup analysis indicates differences in HAM10000 by skin tone (balanced-accuracy gap of 0.176) but the numbers of participants are low, the ITA is indirect and there are few images of darker skin included in HAM10000, and thus no claims can be made about fairness. The Grad-CAM++ maps for nevi and dermatofibroma, which indicate areas beyond the lesion, are also indicative of shortcut learning, thus further accounting for poor transfer.

### 5.7 Efficiency

The proposed model is on the larger size (31.4 M parameters, 5.6 GMACs, 126 MB) and on a single CPU thread, it costs 383 ms per image, in contrast to a few milliseconds on GPU. Thus, it may not be optimal for mobile deployment but is better suited for server-side or workstation inference. The results of static INT8 quantisation (accuracy 0.483) indicate that there is still room for the quantisation procedure to be improved before deployment on the edge: for instance by using quantisation-aware training, or by keeping attention layers in higher precision. In comparison, the MobileNetV3-CBAM baseline (4.3 M parameters, accuracy 0.833) provides a much lighter alternative at the expense of 3.6 accuracy points and 0.09 balanced accuracy.

### 5.8 Segmentation

Networks with pretrained encoders were consistently superior to a standard U-Net trained from scratch in the separate segmentation benchmark (ΔDice 0.012–0.015, all paired tests significant), but the difference between the various pretrained networks was at most 0.003. In our experiments, the 3.1 M-parameter FasterNet-T0 U-Net is therefore the optimal accuracy-cost balance (Dice 0.901, 0.96 GMACs, 5.3 ms), which is oriented towards applications in point-of-care scenarios.[17, 18] The Attention U-Net was not consistently helpful, and the result in the classification ablation was similar, as the Attention U-Net did not outperform the plain FasterNet-T0 U-Net. There are two caveats to the comparison with the standard U-Net: First, the standard U-Net was not pretrained, and second, the validation Dice was still improving at the max epoch. The segmentation and classification experiments were performed on separate datasets and not used together and therefore this study does not demonstrate that lesions masks help with diagnosis. This is still an open question which fuels the joint models presented in Sec. 2.4.

### 5.9 Limitations

All results are based on a single random seed and split, and the effect seen at each split between variants and baselines is a trend and not a proven improvement. (ii) Baselines were trained on a recipe that was different from the proposed model. (iii) HAM10000 has a single-source, lesion-biased case mix and few darker-skin images, and no patient identifier is available, so lesion-wise splitting is the best available safeguard against leakage. (iv) There is an approximate class mapping and two classes have been omitted in the external evaluation. (v) No explanations could be obtained using SHAP for low sensitivity in melanomas. The segmentation experiments are performed using a random image-level split since ISIC 2018 Task 1 doesn’t provide any lesion ID and near duplicate images of one lesion can be present in the training and test sets, so the Dice values reported may be optimistic; further, they are trained using a single seed, and the input size, batch size and learning rate differ among networks, and the training history of Attention U-Net is logged and suggests that training was not performed from scratch and therefore that model should be retrained before submission. (vii) This study is retrospective and has not been clinically validated and the model should not be used for diagnosis.

### 5.10 Future Work

Future plans involve running the experiments multiple times with different seeds and multiple paired statistical tests (McNemar’s for classification / Wilcoxon’s for segmentation), training with domain-robust augmentation, training with data from the smart phone sources, cost sensitive learning to increase melanoma recall, quantisation aware training for deployment, re-analysis of the attention maps to check for shortcut learning, and using the lesion masks created by the model as spatial priors for the classifier in a joint segmentation-classification model.

## 6 Conclusion

This study proposed a hybrid network MaxCAT-Net, consisting of MaxViT-Tiny and CBAM, and a lightweight Transformer encoder for seven-class skin lesion classification, and tested under the leakage controlled (lesion-wise) protocol on HAM10000. The model further outperformed ResNet152-CBAM, MobileNetV3-CBAM, GhostNet-hybrid and single MaxViT-Tiny-CBAM baselines with an accuracy of 0.869, a balanced accuracy of 0.742, macro-F1 of 0.751 and macro-AUC of 0.944. The ablation revealed some minor but consistent improvements with the MaxViT backbone and both CBAM and the Transformer, but not with the GhostNet backbone; the results were not statistically significant given the use of a single seed. Prediction were more reliable when using temperature scaling and deferral based on confidence, as ECE decreased to 0.038 and 94.5% accuracy at 80% coverage. But, when tested on the external validation PAD-UFES-20, all the models tested suffered significant drops in performance, sensitivity for melanoma remained below 0.60 and INT8 quantisation did not work. The 3.1M-parameter FasterNet-T0 U-Net (Dice 0.901) achieved comparable performance to the Swin-T-encoder U-Net (0.903), with only 1/10 of the parameters, and outperformed a standard U-Net trained from scratch (0.888) by 0.02 in the separate segmentation benchmark (ISIC 2018 Task 1). Future efforts are needed towards multi-seed confirmation, cross domain robustness, melanoma-specific training objectives and compression for deployment to clinical setting.

## Disclosures

### Declaration of generative AI use

During the preparation of this work the authors used Claude (Anthropic) for drafting, language editing and the formatting of tables and figures. After using this tool, the authors reviewed and edited the content as needed and take full responsibility for the content of the publication.

### Ethics statement

This study used only existing, publicly released and de-identified image datasets and did not involve the collection of new data from human participants. [Authors: confirm with the institution whether a formal ethics waiver statement is required by the target journal.]

## Code, Data, and Materials Availability

All datasets used in this study are publicly available: HAM10000,[4] ISIC 2018 Task 1,[5] and PAD-UFES-20.[26] The lesion-wise split, the list of removed duplicates and the per-image predictions will be made available at [repository URL to be added]. The training and evaluation code (PyTorch and timm[36]) will be made available at [repository URL to be added].

## Author Contributions (CRediT)

Sadek Al Prince: Conceptualization, Methodology, Software, Validation, Formal analysis, Investigation, Data curation, Visualization, Writing—original draft, Writing—review & editing, Supervision, Project administration. Israt Fatema Shorna: Methodology, Software, Validation, Investigation, Data curation, Writing—original draft, Writing— review & editing. Sadek Al Prince and Israt Fatema Shorna contributed equally to this work.

## Acknowledgments

The authors thank the creators of the HAM10000, ISIC 2018 and PAD-UFES-20 datasets for making their data publicly available.

